# Environmental modifications of dung beetle larvae shape their growth and life history

**DOI:** 10.1101/2025.06.10.658956

**Authors:** Nathan McConnell, Patrick Rohner

## Abstract

Organisms are not just passive recipients of environmental pressures but are able to shape the environment they experience. Yet, the mechanisms and the evolutionary implications of such niche construction remain poorly understood. Here, we study these effects in the gazelle dung beetle (*Digitonthophagus gazella*). Larvae of this species develop in an underground brood chamber (a so-called ‘brood ball’) consisting of cow dung which serves as a sole source of food for a single developing larva. Throughout its development, the larva extensively modifies its environment by constantly eating, regurgitating, and shaping particle sizes within the brood ball. Previous research suggests that these larval manipulations increase environmental quality and nutrient availability. However, how larval modifications affect larval growth and how these modifications differ between species remain poorly understood. We studied the impact of larval environmental modifications by transplanting eggs into previously modified or unmodified environments, whilst controlling for the confounding effect of maternally derived microbes. Additionally, we also studied how *D. gazella* larvae grow in an environment that was modified by a different species (*Onthophagus binodis)* to investigate species-specific differences of niche construction. Counter to expectations, we found that larval modifications by conspecifics did not confer a fitness benefit to *D. gazella*. However, surprisingly, individuals developing in a brood ball that was modified by a heterospecific individual emerged significantly quicker. These findings thus provide mixed support for the hypothesis that environmental modifications by a larva enhance its growth. Our research adds to the growing literature on the complex interactions between organisms and their environment and how those interactions feedback on organismal development and performance.

## Introduction

Natural selection acts on the fit between an organism’s phenotype and its environment. Understanding the relationship between organisms and their environment is thus fundamental to evolutionary biology and there is a wealth of literature documenting how the environment drives (adaptive) evolution (Wallace 1877; Endler 1977). However, while organisms are often seen as passive recipients of environmental selective pressures, many actively shape the way they interact with their surroundings in diverse ways. For instance, organisms can change their phenotype in response to environmental conditions through developmental plasticity. This allows organisms to track changing fitness optima due to changes in the environment (DeWitt et al., 2004; West-Eberhard, 2005; Ghalambor et al., 2007). Furthermore, many organisms can shape the environment they experience through behavioral or physiological mechanisms that directly modify their environment, a process known as niche construction (Odling-Smee et al. 2003; Matthews et al. 2014). By changing the environment an organism experiences, niche construction can in turn mediate plastic responses of the organisms in the modified environment. Understanding how an organism shapes its environment and responds to new environmental conditions is thus a central question in evolutionary biology. We here study these effects in dung beetles.

Dung beetles are an important group of detritivores that provide vital ecosystem services (Krell et al., 2024; Anderson et al. 2024) and are particularly important in agricultural grasslands (Losey et al., 2006; Beynon et al., 2015). In tunneling dung beetles, females (sometimes with the help of a male) bury a dung mass, meticulously shape it into a so-called “brood ball,” and typically deposit a single egg (Hanski et al., 1991). Once the larva hatches, it feeds on the brood ball and rapidly increases its weight about 25-fold before undergoing complete metamorphosis into an adult. Because the larvae spend their entire juvenile period inside their brood ball feeding on their recalcitrant diet, the properties of the brood ball have major implications for larval growth, as well as adult life history and fitness. For instance, it is well documented that maternal provisioning in terms of brood ball size, quality, and burying depth underground affects offspring development time, adult size, and survival (Shafiei et al., 2001; Carter et al., 2020; Kirkpatrick et al., 2022). However, larvae are not solely dependent on maternal provisioning but also actively modify their developmental environment. Immediately after hatching, they begin feeding on and restructuring the contents of their brood ball. Using their heavily sclerotized mandibles, larvae chew and digest the fibrous material, excrete into the brood ball, and repeatedly ingest the altered mixture. Over the course of development, they thus work their way through the brood ball multiple times and heavily modify the mixture until they reach the pupal stage and undergo metamorphosis. At that stage, the brood ball has been transformed into a much finer and more processed material than the original cow dung provided by the mother (Estes et al., 2013; Schwab et al., 2017, Rohner et al., 2024).

Past experiments show that preventing larvae from physically modifying their brood ball results in longer development time and smaller adult size, suggesting that the physical modifications increase environmental quality and nutrient availability (Rohner et al, 2024b). The mechanisms underlying this are becoming increasingly well understood. Previous research shows that larval development is affected by the presence of microbes and that larval modifications enrich microbial communities with taxa able to digest complex carbohydrates (Schwab et al., 2017). Larval behavior might thus help establish an ‘external rumen’ which leverages the establishment of microbial symbionts to help predigesting otherwise hard-to-digest food sources (as is the case in other systems: Flint et al., 2008; Brune, 2014; Béchade et al., 2022). By physically modifying its brood ball and repeatedly digesting the same material, the developing dung beetle might shape its environment in such a way that the nutritional resources within it become more accessible. This could explain why larvae that are repeatedly placed into a new brood ball, and therefore is exposed to much more fresh dung, grow slower and emerge later than larvae that have permanent access to a much smaller quantity of cow dung (Schwab et al., 2017).

Past research suggests that beneficial microbes contribute to beetle growth and life history (Schwab et al. 2016). Intriguingly, part of the microbiome is vertically inherited from mother to offspring (sometimes referred to as the maternal gift) and these maternally inherited microbes have been shown to benefit larval growth (Rohner et al, 2024b, Schwab et al., 2016). These ecologically inherited relationships are thus thought to play a major role in larval development. However, larvae that do not receive a microbial inoculate, while growing slower, do not show higher mortality and undergo otherwise normative development, implying that larval development is not obligately dependent on maternal microbes (Rohner et al, 2024b) (unless exposed to severe environmental stress: see Schwab et al 2016). Recent work by Jones et al., (2024) also shows that, while the maternal microbial inoculant is similar to that of cow dung and the egg, the larval gut microbiome diverges from these sources and is much more similar to the pupal microbiome. This suggests that maternally inherited microbes might play a primary role early in development but that their effect might decrease rapidly over the course of larval development. In addition, the effect size of the presence of maternal microbes is substantially smaller than that of a larva’s ability to physically modify its environment (Rohner & Moczek 2024). This suggests that the actions of the larva itself have substantial effects on development and might ultimately have a stronger effect on development time and body size. However, how these physical modifications affect larval growth is poorly understood.

To better understand how larval environmental modifications influence development, we study the dung beetle *Digitonthophagus gazella*. Specifically, we investigate whether physical alterations to brood balls by larvae feedback to enhance survival, growth rate, and adult size. By explicitly excluding maternally derived microbes, we isolate the effects of larval-driven modifications. To assess whether such modifications are adaptive, we introduce newly hatched larvae into standardized artificial brood balls (ABBs) previously modified by conspecifics and compare their performance to larvae placed in unmodified artificial brood balls. To explore the species-specificity of these modifications, we also expose *D. gazella* larvae to artificial brood balls previously modified by a closely related species, *Onthophagus binodis*. If larval modifications have evolved to match species-specific requirements, we expect larvae to perform best in environments modified by conspecifics. Additionally, we include two control treatments to distinguish the effects of larval modification from those of unmodified microbial communities. Taken together, our results indicate complex and unexpected relationships between larval niche construction and developmental outcomes.

## Methods

### General laboratory populations

We focus on the gazelle dung beetle, *Digitonthophagus gazella,* originally collected in Bastrop County, Texas. For comparison we also used *Onthophagus binodis,* originally from Waimea, Hawaii. These two species diverged about 40 million years ago (Parzer et al., 2018) but are ecologically similar. Both species are native to sub-Saharan Africa and overlap in their native and non-native ranges (Tyndale-Biscoe, 1990; Davis et al., 2020). Whilst *D. gazella* is found on a wider range of animal dung, both species are commonly found on cattle dung (Davis et al., 2020). *O. binodis* is smaller in adult size compared to *D. gazella*, however, the mean larval body weights over the first eight days fall within a comparable size range (Rohner et al., 2021). Both species were maintained in the laboratory in large colony containers filled with substrate (a previously autoclaved mixture of sand and topsoil) and fed with defrosted cow dung once a week.

### Experimental design

To expose focal larvae to environments that varied in the degree to which they were modified by a larva, we conducted an experiment with two distinct phases. In Phase I (Fig.1) we generated experimental artificial brood balls (ABBs) that differed in the degree to which they were modified by larvae, including modification by a conspecific *D. gazella*, a heterospecific *O. binodis,* as well as two control treatments that accounted for potential effects that are driven by the age of an unmodified brood ball. In Phase II, (Fig. 2) we exposed focal individuals to these experimental artificial brood balls and quantified how the presence (or absence) of larval manipulations affected larval growth. Below, we first describe how we generate the treatments in Phase I and how we then assessed their effect on beetle growth during Phase II.

**Fig. 1.**
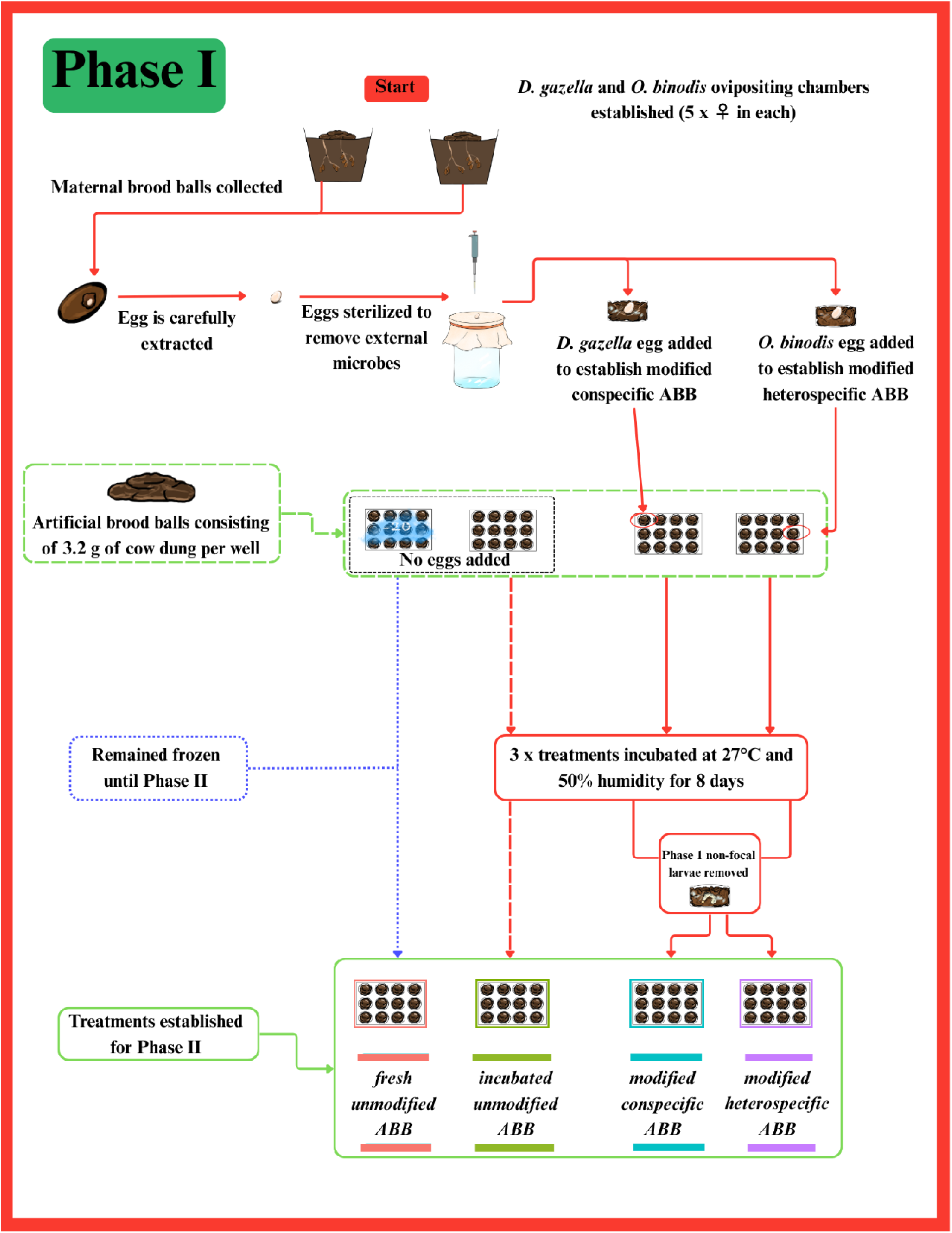
Flow chart describing the process of generating the artificial brood ball treatments treatments in phase I.

**Fig. 2.**
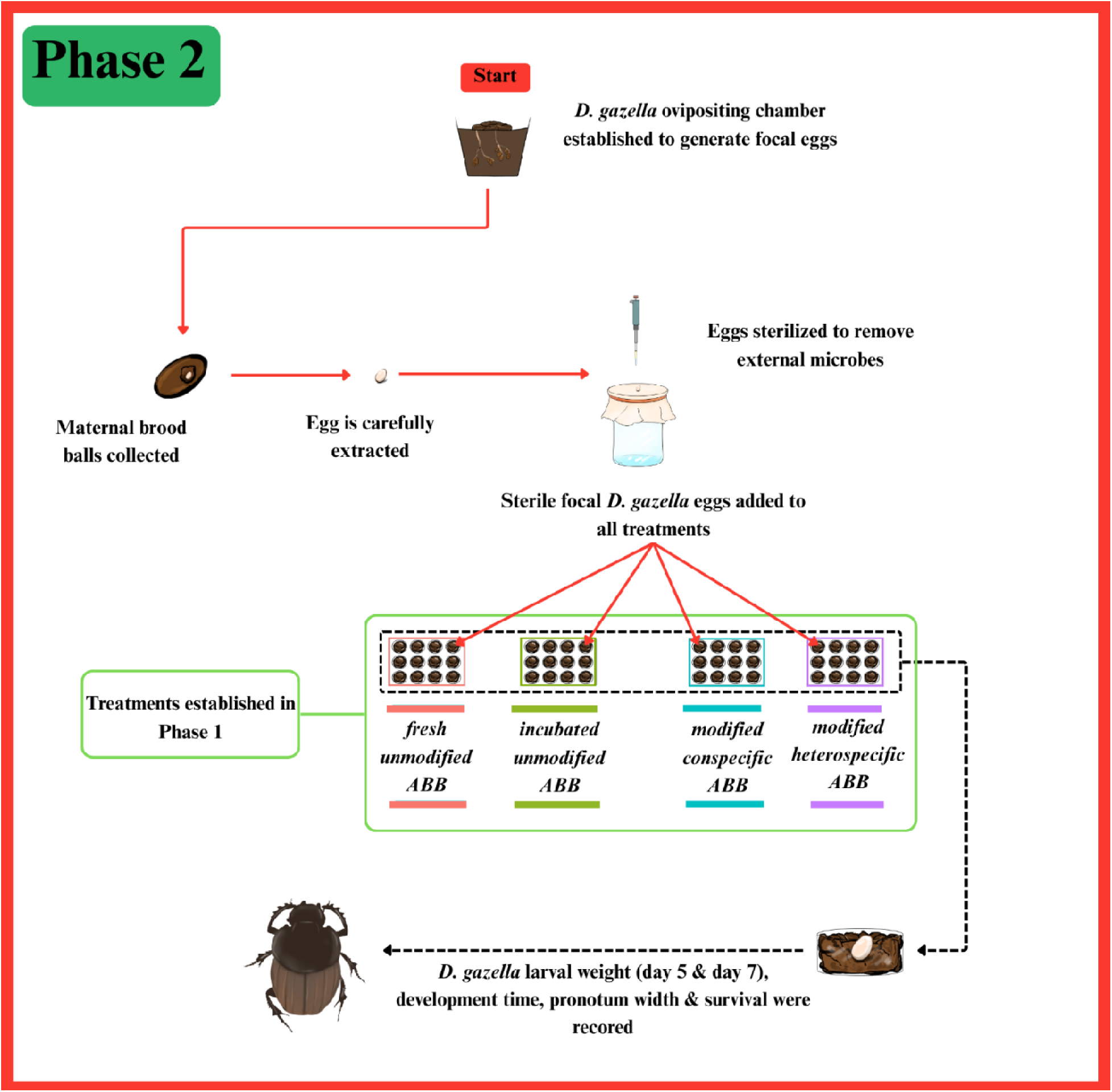
Flow chart describing phase II.

### Phase I: generating experimental brood ball treatments (*Fig 1*)

To generate experimental brood balls, we first made ‘artificial brood balls’ (ABB; following standard procedures: Shafiei et al., 2001). In brief, previously frozen cow manure was squeezed by hand in a cheese cloth, removing excess moisture to replicate the consistency of natural brood balls. 3.2g of this manually processed dung was then placed in the wells of sterile 12-well plates (VWR 734-2324). Each well then represents a single artificial brood ball. To limit experimental noise, all brood balls were made from the same batch of defrosted cow dung and frozen. Note that, although freezing eliminates invertebrates, it preserves a diverse microbial community (Ledón-Rettig et al., 2018). We established two control treatments and two experimental brood ball treatments differing in larval modification history:

1. *fresh unmodified ABB*: artificial brood ball kept frozen during Phase I and defrosted just before Phase II.
2. *incubated unmodified ABB*: artificial brood ball incubated for 8 days without a larva.
3. *modified conspecific ABB*: artificial brood ball modified for 8 days by a *D. gazella* larva
4. *modified heterospecific ABB*: artificial brood ball modified for 8 days by an *O. binodis* larva.

To generate the larvae needed to establish the treatments we removed ten mature females of *D. gazella* and *O. binodis* from the general populations and placed them in two circular 235mm x 210mm ovipositing chambers. The ovipositing chambers contained a compact mixture of sterilized soil and sand and were topped with previously frozen cow dung (approx. 250g) and were kept in a controlled insect rearing chamber (Caron) at 27°C and 50% humidity for six days. After six days, all maternal brood balls were carefully extracted by sifting the soil from the ovipositing chamber. The maternal brood balls were carefully opened, and eggs were extracted from the maternal brood ball using flame-sterilized featherweight forceps and placed on a sterilized gauze mesh. Each egg was then washed with 500µl of sterilization liquid (48.5ml dH20, 0.5ml bleach, 0.05ml Triton X; see Rohner et al., 2024b; Schwab et al., 2016) followed by two rinses of distilled water (2 x 500µl) to eliminate the potentially confounding effects of maternal microbiomes. Sterilized eggs were placed into newly defrosted artificial brood balls. Only one egg of either *D. gazella* or *O. binodis* was allocated to each artificial brood ball, establishing the *modified conspecific* and *modified heterospecific ABB* treatments, respectively. The lids of the 12-well plate were closed and transferred into an incubator (Caron Insect Rearing Chamber) set at 27°C and 50% humidity.

To establish control treatments that lacked larval modifications we used two different approaches. Firstly, we kept a set of artificial brood balls frozen for the duration of Phase I. This ensured that, at the start of Phase II, we had a set of artificial brood balls that replicate a fresh unmodified environment (*fresh unmodified ABB)*. Secondly, we also incubated artificial brood balls for 8 days in the same incubator as the modified treatments but without a larva (*incubated unmodified ABB)*. This treatment thus lacked larval manipulations but allows for any potential microbial growth or other environmental changes that accrue over time in the absence of larval activity.

At the end of day 8 of Phase I, we removed all non-focal larvae from their brood balls (for those two treatments that had larvae). Brood balls, within which eggs did not hatch or larvae did not survive until day 8 were excluded from the experiment.

### Phase II: quantifying impacts on larval growth (*Fig 2*)

Two days into Phase I, four *D. gazella* ovipositing chambers were set up to generate focal individuals after six days. Focal eggs were extracted from the maternal brood balls, washed with fresh sterilization liquid, and placed into the four different brood ball treatments that were generated in Phase I. All plates containing artificial brood balls were closed with a lid and transferred to an incubator (Caron Insect Rearing Chamber) set at 27°C and 50% humidity. Due to the relatively long development and low fecundity of dung beetles and logistical constraints, the experiment was repeated in three consecutive temporal blocks over one month, totaling 161 individuals that hatched from 217 eggs.

Eggs were checked daily to assess the potential effect of the brood ball treatment on egg hatching success and to control the age of larvae later in the experiment. To quantify larval growth rate, we recorded total larval body weight on day five and day seven post hatching for all surviving larvae. Larval weight was measured using a Mettler Toledo scale (Model AB135-S) to three decimal places. Insect growth curves are intrinsically nonlinear and often complex. We therefore focused on the initial “free” period of larval growth which approximates the maximal physiologically realized growth rate (Rohner et al., 2021). As an estimate of larval growth, we calculated instantaneous relative growth rates (RGR) following (Tammaru et al., 2007):

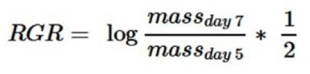

This estimate better approximates larval growth dynamics in a growing larva than “integral” estimates of growth rate that divide adult size by egg-to-adult development time (Tammaru et al., 2007).

Daily mortality checks were conducted visually until all individuals had emerged from the pupal stage, and adults were collected three days after emergence to allow cuticular hardening. The widest basal point of the pronotum was used as a proxy for body size, as established in dung beetle literature (Rohner et al., 2023). Pictures of the pronotum were obtained with a Pixelink camera (M20C-CYL) attached to a LEICA M205 microscope and measured in ImageJ (Version 1.54m) (Schneider et al., 2012).

### Statistical analysis

All statistical analysis was performed using R V-4.4.1 (R Core Team 2024) (Code provided in supplementary material). Egg hatching success and larval survival were recorded as binary variables and were consequently analyzed using generalized mixed effects models from the lme4 package with a binomial link function (Bates et al., 2015). The three separate temporal replicate blocks were analyzed simultaneously, with experimental block designated as a random effect. Treatment was used as the only fixed effect.

Development time, relative growth rate (RGR) and pronotum width (adult size) were analyzed using mixed effect models from the lme4 package (Bates et al., 2015). Block was again added as a random effect. Treatment and sex were used as fixed effects. We always fitted full models, including the sex-by-treatment interaction, but excluded the interaction term in cases where it was statistically not significant (following standard backward elimination procedures: Neter et al 1985). For all models, Q-Q plots and histograms were used to check the residual distribution. A singular individual with an unusually large RGR had strong impacts on the analysis and residual distribution and was therefore removed from the analysis. Where appropriate, differences between group treatments were analyzed using a Tukey post hoc test from the ‘emmeans’ package (Lenth, 2024) or from the ‘multcomp’ package (Hothorn et al., 2008). Effect size was calculated using partial eta-squared (η_p_^2^) values from the package ‘effectsize’ (Ben Shachar et al., 2020).

## Results

We assessed whether larval modifications to brood balls feed back on larval growth and performance by exposing developing larvae to brood balls modified in various ways. Neither egg hatching rates (Fig. S1) nor larval survival (Fig. S2) differed among the four treatments (hatching success: χ² = 7.01, df = 3, p = 0.072, n = 217; larval survival: χ² = 4.90, df = 3, p = 0.179, n = 161), suggesting that brood ball manipulations do not affect mortality during early life stage transitions.

However, brood ball treatment significantly affected egg-to-adult development time (χ² = 50.05, df = 3, p < 0.001, η_p_^2^= 0.50, 95% CI: [0.27, 0.64]; Fig.3). Specifically, larvae developing in a brood ball previously modified by a conspecific took nearly five days longer to reach adulthood than larvae developing in fresh unmodified artificial brood balls (estimate = 4.93 ± 1.00 days, p < 0.001). Contrary to expectation, larvae inhabiting a brood ball that was manipulated by a larvae of the same species tended to take longer to develop compared to those in brood balls manipulated by a *different* species, although this effect was marginally non-significant (Tukey-adjusted comparisons between modified *conspecific vs. heterospecific ABBs* = 3.01 ± 1.16 days, p = 0.057). In addition, larvae inhabiting a fresh unmodified brood ball had significantly shorter development times compared to larvae placed in brood balls that were left empty for 8 days (difference between *fresh vs. incubated unmodified ABBs*: estimate = 5.46 ± 1.03 days, p < 0.001). We found no significant effects of sex (χ² = 0.04, df = 1, p = 0.841) or evidence of an interaction between treatment and sex on development time (χ² = 3.57, df = 3, p = 0.312).

**Fig. 3.**
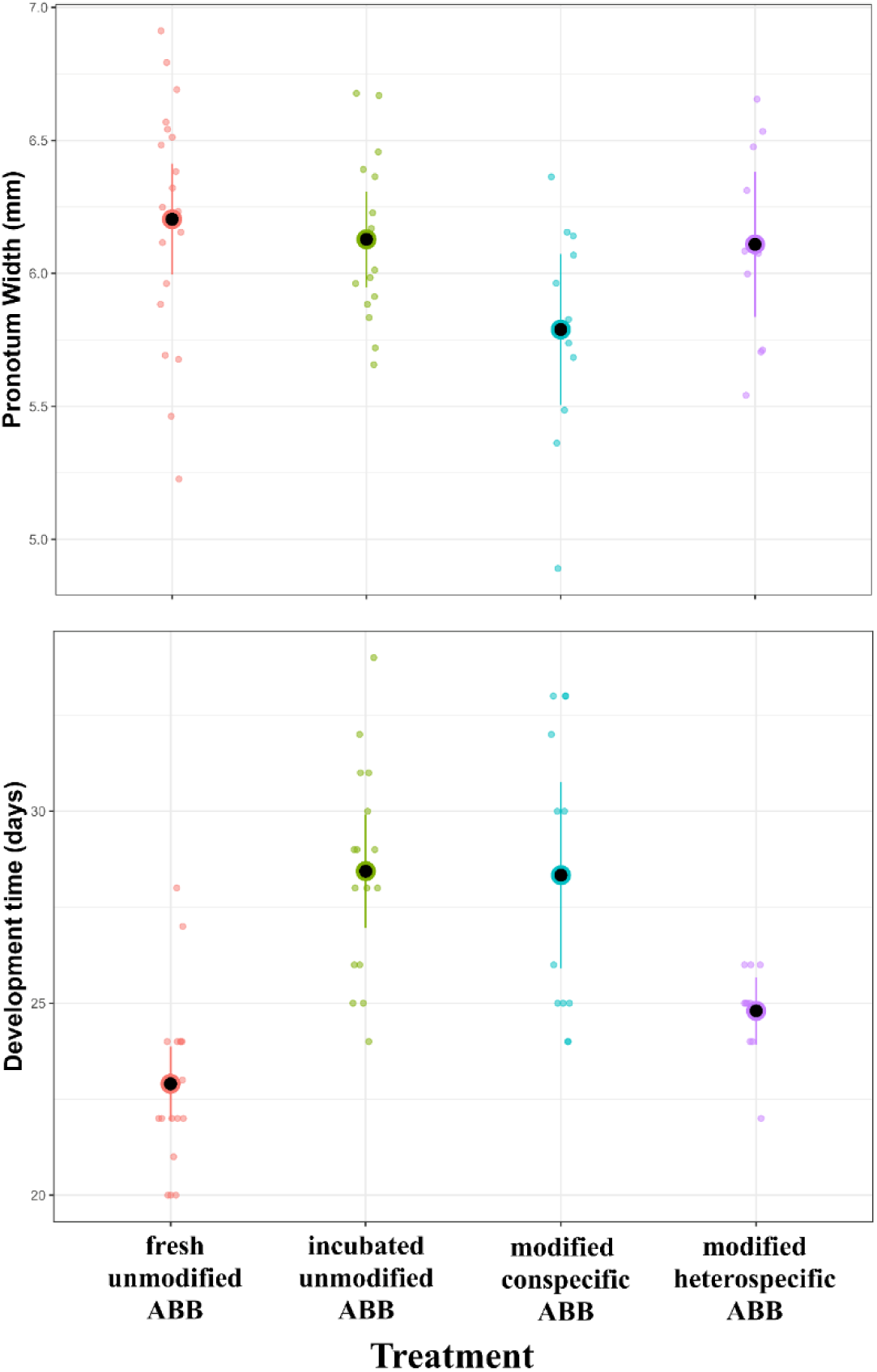
Egg to adult development time and adult body size (pronotum width) for larvae reared in different artificial brood ball (ABB) treatments. Plots show means (Black dots) and corresponding 95% confidence limits. Colored points indicate treatment-specific individuals.

Larval weight at day five was affected by treatment (χ² = 52.68, df = 3, p = < 0.001, η_p_^2^= 0.50, 95% CI: [0.29, 0.64] ; Fig.4), but not sex (χ² = 0.22, df = 1, p = 0.641). Pairwise comparisons show that individuals in a fresh unmodified artificial brood ball were heavier than larvae in all other treatments (all p < 0.019).

**Fig. 4.**
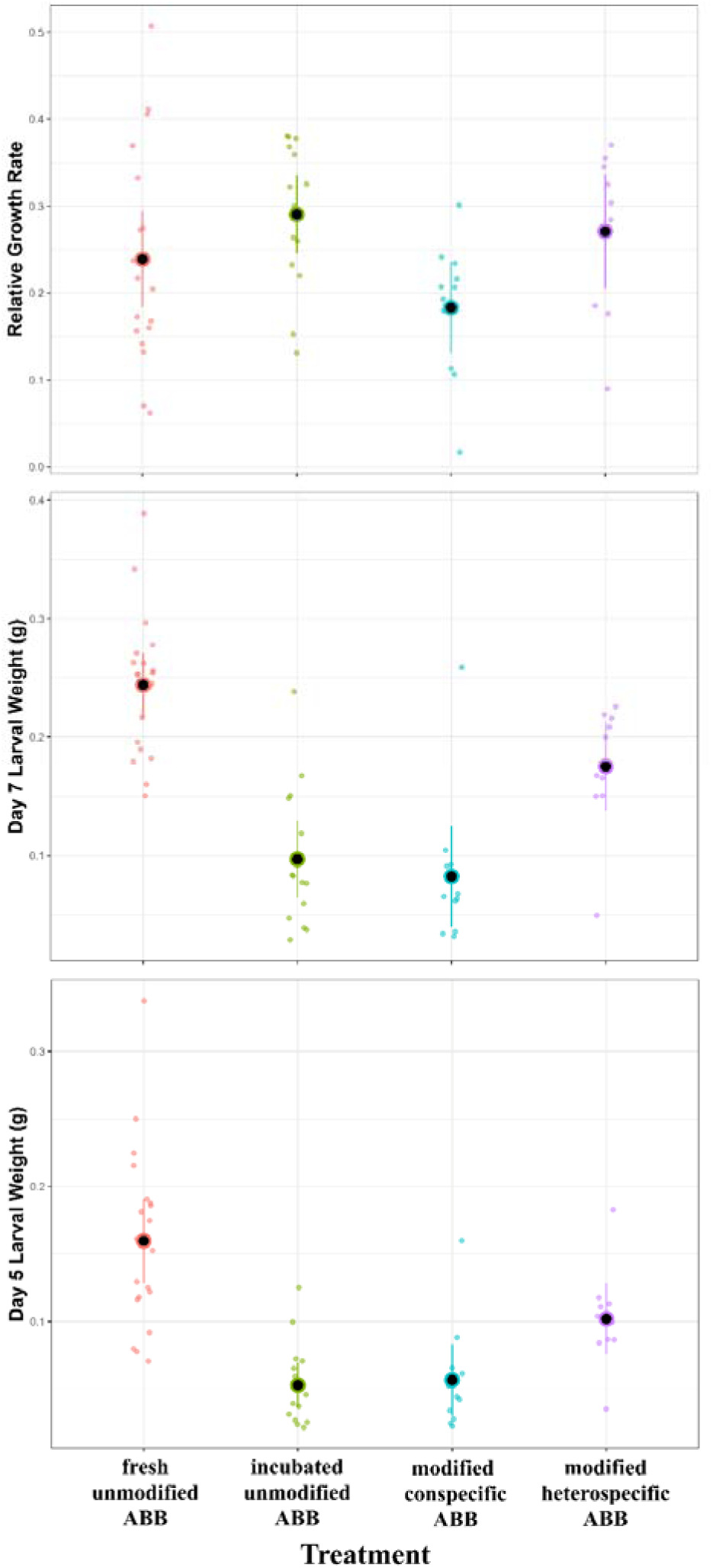
Relative growth rate and Larval weight at day 5 and day 7 for larvae reared in different artificial brood ball (ABB) treatments. Plots show means (Black dots) and corresponding 95% confidence limits. Colored points indicate treatment-specific individuals.

Larval weight at day seven followed a similar trend to that at day five, with a significant effect of treatment (χ² = 76.57, df = 3, p = <0.001, η_p_^2^= 0.68, 95% CI: [0.51, 1.00] ; Fig.4), but not sex (χ² = 0.04, df = 1, p = 0.836). Larvae in fresh unmodified brood balls were significantly larger than larvae in the other treatments (all pairwise differences p < 0.018). Interestingly, the pairwise comparisons also show that focal larvae developing in brood balls that were manipulated by a different species were larger than larvae in brood balls manipulated by a conspecific (*modified heterospecific vs. conspecific ABBs:* estimate = 0.091 ± 0.026, p = 0.005). Larvae developing in a modified heterospecific artificial brood ball were also larger than those inhabiting incubated unmodified brood balls (estimate = 0.078 ± 0.026, p = 0.025). This further emphasizes the influence that the early environment has on dung beetle development time.

Relative larval growth rate showed a significant effect of treatment (χ² = 7.84, df = 3, p= 0.049; Fig.4). However, effect sizes were relatively low (η_p_^2^= 0.14, 95% CI: [0.00, 0.30]) and pairwise comparisons were not significant. Pronotum width (Fig. 3) as a measure of adult body size was again significantly different by treatment (χ² = 7.99, df = 3, p= 0.046), but not sex (χ² = 2.14, df = 1, p= 0.144) or interaction between treatment and sex (χ² = 2.49, df = 3, p = 0.477). Pairwise post hoc comparisons reveal that individuals that developed in fresh unmodified brood balls were larger than individuals in brood balls that were modified by a conspecific (*modified conspecific ABBs*: estimate = 0.407 ± 0.15, p = 0.043). As with relative growth rate, the effect size was modest (η_p_^2^= 0.17, 95% CI: [0.00, 0.35]). Collectively, these results show that larval brood ball modifications do not universally enhance larval growth. Instead, the effect of larval modifications emerges as context-dependent and only partially consistent with the hypothesis that larval modifications enhance larval growth.

## Discussion

Dung beetles, like many other organisms that digest complex carbohydrates, are thought to depend on environmental modifications to extract nutritional value from nutritionally poor or hard-to-digest substrates (Hungate, 1984; Parker et al., 2019). Here, we tested whether larval manipulation of the brood ball improves larval development and whether the extent of this benefit is species-specific. In contrast to our expectations, we found little evidence of beneficial modifications to the brood ball. Irrespective of which variable measured, exposing *D. gazella* larvae to a previously modified brood ball did not improve developmental outcomes compared to previously unmodified brood balls. Although our experimental approach controlled for the potentially confounding effect of maternal microbes, this suggests that larval modifications do not necessarily improve larval development. Instead, we recovered an unexpected pattern across species, suggesting more complex ecological interactions between larvae and their ontogenetic environment that are not fully consistent with a conserved adaptation.

### Larval brood ball modifications do not necessarily benefit larval growth in D. gazella

Dung beetle larvae are known to drastically restructure and modify their brood ball throughout their juvenile development (Hanski et al., 1991, Holter, 2016 and Madzivhe et al., 2021). These modifications are thought to construct a more suitable ontogenetic environment, including the establishment of microbial communities that are able to digest complex compounds (Schwab et al., 2017, Chen et al., 2024), specific to their life stage (Shukla et al., 2016). These previous observations are thus consistent with the idea that dung beetle larvae establish an external rumen. However, if larval modifications were to establish an external rumen, we expected larvae to grow the fastest when placed in brood balls that had previously been inhabited by another conspecific larvae (*modified conspecific ABB*). In contrast, the growth of larvae in modified conspecific ABBs were indistinguishable from the growth of larvae from incubated or fresh unmodified ABBs. Surprisingly, larvae placed in fresh unmodified ABBs were heavier than larvae in any other treatment at days 5 and 7 and they developed faster than larvae placed in brood balls that were previously modified. This implies that larval modifications do not necessarily benefit larval growth. This also demonstrates that the early larval environment has a more significant impact on development time than any accrual of beneficial modifications to the microbiome that may occur later. For example, the advantage of inhabiting a newly defrosted (*fresh unmodified ABB*) over an old *(incubated unmodified ABB)* one is to emerge as an adult almost five days earlier, and that is seemingly correlated with the larval weight established in those initial first few days. Taken together, these findings suggest larval niche construction in *D. gazella* to be at best ineffectual, whilst in some instances detrimental, and ultimately, inconsistent with the idea that larval modifications generally benefit larval growth.

### Potential evolution of larval brood ball manipulation

Organism-environment interactions are taxon-specific (Holt et al., 2024; Schwab et al., 2017) and, if adaptive, one would expect larvae to perform best in a brood ball that was modified by a conspecific. However, surprisingly, *D. gazella* larvae emerging from a brood ball that was modified by a *O. binodis* larva were heavier at day 7 compared to those emerging from brood balls previously inhabited by a conspecific. This indicates that environmental manipulation can be beneficial, at least in some circumstances, and that these effects diverge between species. The mechanisms underpinning these species-specific effects are, however, unclear. For instance, *O. binodis* may be more capable of constructing a beneficial niche than *D. gazella*. Alternatively, these effects could be explained by the initial *D. gazella* consuming a significantly larger proportion of the brood ball, thus, leaving the second (focal) larva with less available resources. It has been demonstrated that poorly provisioned brood balls directly impact the size of emerging dung beetles (Shafiei et al., 2001). However, the initial artificial brood ball was provisioned with 3.2g of dung, which is much more than required to reach maximum size (Rohner et al., 2021). Additionally, the eight-day rearing standard was determined by intersecting larval weight between both species (Rohner et al., 2021), whilst maximizing the time window for larval modifications to accrue. Visual inspection of the non-focal *D. gazella* larvae used to establish the modified brood ball in Phase I did appear larger than *O. binodis* larvae after eight days. However, the amount of dung available to the newly introduced focal *D. gazella* egg in *modified conspecific ABBs* did not seem significantly deficient, though it was more fibrous than *modified heterospecific ABBs* (note that all treatments were maintained under 27°C and 50% humidity). The observed differences may indicate that aspects of larval brood ball modifications evolve. On one hand, *O. binodis* might simply be better modifiers of their brood ball than *D. gazella*. Alternatively, the two species might differ in their target resources in early development, resulting in a depletion of specific compounds established within the brood balls of a conspecific versus a heterospecific. Future work will be necessary to uncover the mechanistic basis of these species differences.

### The potential role of vertically transmitted microbes

It is important to consider that we here focus exclusively on the impact of larval behavior on brood ball modification while explicitly excluding parental effects, especially the maternally inherited microbiome (maternal gift). Previous studies indicate that these factors are interacting in complex ways. The presence or absence of a maternal pedestal significantly affects the development time of females, but only when larval modifications to a brood ball are disrupted, indicating that there are sex specific effects on the disruption of maternal microbes (Rohner et al., 2023; Rohner et al., 2024a). To fully reject the hypothesis that larval modifications themselves are adaptive, future research will also need to jointly manipulate the presence of maternal microbiomes and larval modifications. This was not feasible in our experimental design because species specific larval modifications would have been experimentally confounded with species-specific differences in vertically-transmitted microbiomes.

However, although we cannot exclude interactions with the maternal microbiome, the lack of evidence that brood ball modifications improve larval performance is still surprising and points to alternative mechanisms. The striking difference between the *fresh unmodified ABB* and *incubated unmodified ABB* on growth outputs suggests that, instead of benefiting the establishment of beneficial microbes that increase nutrient availability, larval modifications might instead suppress the growth of harmful microbial taxa. *Incubated unmodified ABB* accumulated visible microbial communities, sometimes leading to a large aggregation of fungal fruiting bodies. This was neither seen in the freshly defrosted control, nor in brood balls that were inhabited by a larva, suggesting that larval activity might disrupt the establishment of particular microbial communities. The suppression of harmful microbes is widespread across taxa (e.g., Amphibians: Harris et al., 2006; Fish: Masso-Silva et al., 2014; Reptiles: Van Hoek, 2014). Previous work on Grahams’s Frog (*Odorrana graham*) has recovered more than 370 antimicrobial peptides produced in its skin (Li et al., 2007) and in birds; the European Hoopoe (*Upupa epops*) uses metabolites produced by a symbiotic bacteria living in their uropygial gland that inhibits the growth of harmful microbes (Martín-Vivaldi et al., 2009). In insects the larvae of the fruit fly *Drosophila melanogaster* suppress the specific filamentous fungi growth through communal aggregations of larvae (Trienens et al., 2020). Whilst both the larvae of coconut rhinoceros beetle (*Oryctes rhinoceros*) and black soldier fly produce antimicrobial peptides effective at suppressing harmful microbes (Yang et al., 1998, Zhang et al., 2022), highlighting a common microbial defense strategy among insects.

Alternatively, the uninhibited microbial growth *incubated unmodified ABBs* may have exhausted the nutritional value of the dung, thus slowing larval development time. However, if nutrition was restricted then we might expect body size to be significantly smaller (Rohner et al., 2021). Therefore, the function of preventing certain microbes from being established thus appears more likely than the depletion of nutrition in the brood ball.

### Conclusions

Previous studies suggested that dung beetle larvae modify their environment in ways that benefit their growth (Schwab et al., 2016; Parker et al., 2019; Rohner et al., 2023, 2024b) and earlier research suggested the existence of an external rumen that increases nutrient availability. In contrast to these expectations, we only found limited support for increased nutrient availability through larval modification in *D. gazella*. Instead, our results suggest that brood ball modifications might hinder the establishment of harmful microbes. More generally, our study highlights the complex and unexpected interplay between developing organisms, their effects on the environment, and how modified environments feed back on organism growth, life history, and fitness (Rohner et al., 2024a; Sultan, 2015, Natta et al., 2024). Rather than viewing the relationship between a dung beetle and their environment as a linear, unidirectional process in which environmental pressures simply shape selective outcomes, our findings suggest a more dynamic and reciprocal interaction. Organisms do not merely respond to environmental challenges; instead, they actively reshape their surroundings, which in turn influences their own development and potentially that of future generations. Future research will be required to shed light on the detailed microbial and behavioral mechanisms mediating these effects in dung beetles.

## Acknowledgements

We would like to thank collaborators who helped with collection, husbandry and stimulating discussions around this manuscript including Ben Mathews, Dr. Michelle Herrera, Dr. Sarah Britton, Ebony Argaez, and Avi Khanna. We are also grateful for helpful and constructive comments by two Anonymous Reviewers.

## Data availability

Raw data and code available at DOI: 10.5061/dryad.z8w9ghxqv

