## Supplementary material for "Environmental modifications of dung beetle larvae shape their growth and life history"

**Supplementary Materials**

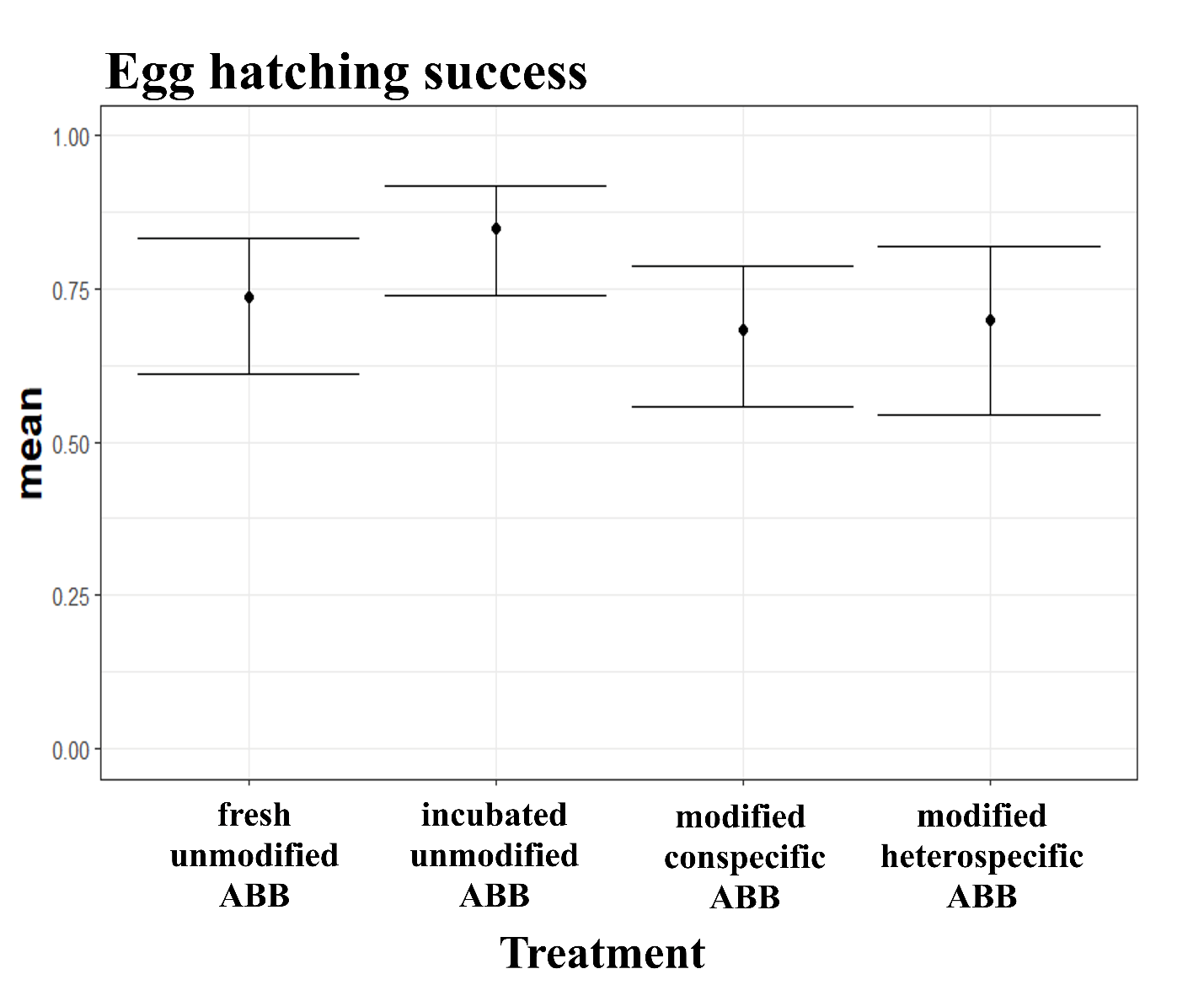

Figure S1: Mean hatching success for experimental treatments; Fresh Unmodified ABB (n = 60), Incubated Unmodified ABB (n = 57), modified conspecific ABB (n= 60) and modified heterospecific ABB (n = 40). Plots show means (Black dots) and corresponding 95% confidence limits.

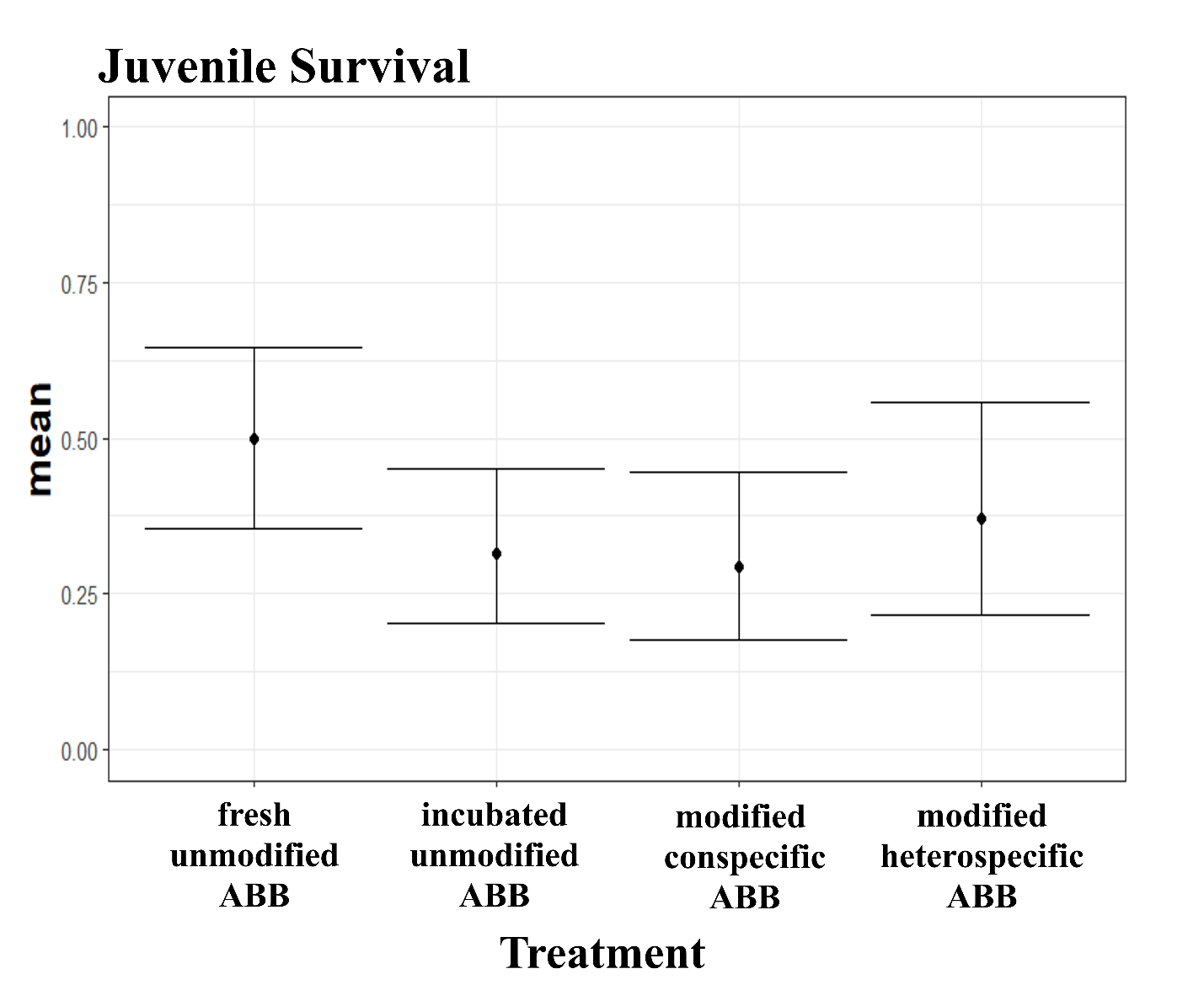

Figure S2: Mean larval survival for experimental treatments; Fresh unmodified ABB, incubated unmodified ABB, modified conspecific ABB and modified heterospecific ABB. Plots show means (Black dots) and corresponding 95% confidence limits.

\# Analysis Code for Manuscript: ["The external rumen of dung beetles:

\# Complex interactions between larvae and their ontogenetic environments

\# shape growth and life history"]

\# Author: [Nathan John McConnell]

\# Date: [3/20/2025]

\# Description: R code for analyzing hatching success, larval survival,

\# development time, growth rate, and pronotum width of *Digitonthophagus gazella*

```
# Required Libraries
library(ggplot2)
library(dplyr)
library(lmerTest)
library(emmeans)
library(effectsize)
library(car)

# Set theme
theme1 <- theme_bw() +
  theme(
    panel.grid.major = element_blank(),
    panel.grid.minor = element_blank(),
    axis.title.x = element_text(size = 16, face = "bold"),
    axis.title.y = element_text(size = 16, face = "bold"),
    axis.text.x = element_text(size = 15, face = "bold"),
    strip.text = element_text(size = 15, face = "bold"),
    legend.position = "none"
  )

###############################
# Hatching Success
d <- read_excel("LRT data.xlsx", sheet = "Eggs")
d$treatment <- factor(d$treatment, levels = c("FreshCon", "Control", "Gazella", "Binodis"))
m1 <- glmer(hatched ~ treatment + (1|Block), family = binomial(), data = d)

# Summary and plots for hatching success
summary(m1)
emmeans(m1, pairwise ~ treatment)
summary_hatching <- ddply(d, .(treatment), summarize, n = length(hatched), hatched = sum(hatched))
summary_hatching <- cbind(summary_hatching, binom.confint(summary_hatching$hatched, summary_hatching$n, method = "wilson")[, c("mean", "lower", "upper")])
ggplot(summary_hatching, aes(x = treatment, y = mean)) +
  geom_point() + geom_errorbar(aes(ymin = lower, ymax = upper)) + ylim(c(0, 1)) + theme1 +
  ggtitle("Egg Hatching Success")

###############################
# Larval Survival
d2 <- read_excel("LRT data.xlsx", sheet = "Survival")
d2$treatment <- factor(d2$treatment, levels = c("FreshCon", "Control", "Gazella", "Binodis"))
m2 <- glmer(survived ~ treatment + (1|Block), family = binomial(), data = d2)

# Summary and plots for larval survival
summary(m2)
emmeans(m2, pairwise ~ treatment)
summary_survival <- ddply(d2, .(treatment), summarize, n = length(survived), survived = sum(survived))
summary_survival <- cbind(summary_survival, binom.confint(summary_survival$survived, summary_survival$n, method = "wilson")[, c("mean", "lower", "upper")])
ggplot(summary_survival, aes(x = treatment, y = mean)) +
  geom_point() + geom_errorbar(aes(ymin = lower, ymax = upper)) + ylim(c(0, 1)) + theme1 +
  ggtitle("Juvenile Survival")

###############################
# Development Time
d3 <- read_excel("LRT data.xlsx", sheet = "Development")
d3$treatment <- factor(d3$treatment, levels = c("FreshCon", "Control", "Gazella", "Binodis"))
d3 <- subset(d3, !is.na(Sex))  # Remove specimens with unknown sex

m3 <- lmer(Devtime_Days ~ treatment * Sex + (1|Block), data = d3)
Anova(m3)  # Removed interaction if not significant
summary(m3)
emmeans(m3, pairwise ~ treatment)
eta_squared(m3, partial = TRUE)
omega_squared(m3)

# Plot development time
ggplot(d3, aes(x = treatment, y = Devtime_Days)) +
  geom_point(alpha = 0.5, aes(color = treatment), position = position_jitterdodge(dodge.width = 0.25)) +
  theme1 +
  stat_summary(fun.data = mean_cl_normal, aes(color = treatment), position = position_dodge(width = 0.25)) +
  stat_summary(fun = mean, aes(color = treatment), geom = "point", shape = 21, fill = "black", size = 4, stroke = 1) +
  labs(x = "Treatment", y = "Development Time (Days)")


###############################

# Relative Growth Rate
d4 <- read_excel("LRT data.xlsx", sheet = "Development")
d4$treatment <- factor(d4$treatment, levels = c("FreshCon", "Control", "Gazella", "Binodis"))
d4 <- subset(d4, !is.na(Sex))  # Remove specimens with unknown sex
d4$RGR <- log(d4$'Day7LW' / d4$'Day5LW') / 2
d4 <- subset(d4, RGR < 0.9)  # Exclude outlier

m4 <- lmer(RGR ~ treatment + (1|Block), data = d4)
Anova(m4)
summary(m4)
emmeans(m4, pairwise ~ treatment)
eta_squared(m4, partial = TRUE)
omega_squared(m4)


# Plot relative growth rate
ggplot(d4, aes(x = treatment, y = RGR)) +
  geom_point(alpha = 0.5, aes(color = treatment), position = position_jitterdodge(dodge.width = 0.25)) +
  stat_summary(fun.data = mean_cl_normal, aes(color = treatment), position = position_dodge(width = 0.25)) +
  stat_summary(fun = mean, aes(color = treatment), geom = "point", shape = 21, fill = "black", size = 4, stroke = 1) +
  labs(x = "Treatment", y = "Relative Growth Rate") +
  theme1

###############################
# Larval weight on day 5
m6<-lmer(Day5LW~treatment +(1|Block),data=d4)
summary(m6)
Anova(m6)
emmeans(m6, pairwise~treatment)
eta_squared(m6, partial = TRUE)

ggplot(d4, aes(x = treatment, y = Day5LW)) +
  geom_point(alpha = 0.5, aes(color = treatment), position = position_jitterdodge(dodge.width = 0.25)) +
  stat_summary(fun.data = mean_cl_normal, aes(color = treatment), position = position_dodge(width = 0.25)) +
  stat_summary(fun = mean, aes(color = treatment), geom = "point", shape = 21, fill = "black", size = 4, stroke = 1) +
  labs(x = "Treatment", y = "Larval weight day 5") +
  theme1

###############################
# Larval weight on day 7
m7<-lmer(Day7LW~treatment +(1|Block),data=d4)
summary(m7)
Anova(m7)
emmeans(m7, pairwise~treatment)
eta_squared(m7, partial = TRUE)

ggplot(d4, aes(x = treatment, y = Day7LW)) +
  geom_point(alpha = 0.5, aes(color = treatment), position = position_jitterdodge(dodge.width = 0.25)) +
  stat_summary(fun.data = mean_cl_normal, aes(color = treatment), position = position_dodge(width = 0.25)) +
  stat_summary(fun = mean, aes(color = treatment), geom = "point", shape = 21, fill = "black", size = 4, stroke = 1) +
  labs(x = "Treatment", y = "Larval weight day 7") +
  theme1

###############################

# Adult Pronotum Width
m5 <- lmer(ProWid ~ treatment + (1|Block), data = d4)
Anova(m5)
emmeans(m5, pairwise ~ treatment)
eta_squared(m5, partial = TRUE)
omega_squared(m5)


# Plot pronotum width
ggplot(d4, aes(x = treatment, y = ProWid)) +
  geom_point(alpha = 0.5, aes(color = treatment), position = position_jitterdodge(dodge.width = 0.25)) +
  stat_summary(fun.data = mean_cl_normal, aes(color = treatment), position = position_dodge(width = 0.25)) +
  stat_summary(fun = mean, aes(color = treatment), geom = "point", shape = 21, fill = "black", size = 4, stroke = 1) +
  labs(x = "Treatment", y = "Pronotum Width (mm)") +
  theme1
```
